# Effectiveness of the University Executive Network-Training Program (NExT-U) on executive functions in university students

**DOI:** 10.64898/2026.07.25.740682

**Authors:** Diego D. Díaz-Guerra, Evelyn Fernández-Castillo, Carlos Ramos-Galarza, Melany de la Torre Pérez, Yelenys González Espinosa, Marena C. de la Hernández-Lugo, Vania Lugones Dapresa, Yunier Broche-Pérez

## Abstract

**Introduction:** Executive functions (EF) are higher-order cognitive processes essential for academic performance in university settings. Although there is extensive research on EF training in children, studies in young adults are scarce, particularly those involving interventions tailored to specific needs.

**Objective:** To evaluate the effect of the University Executive Network-Training Program (NExT-U), based on specific needs, on the executive functioning of Cuban university students.

**Methodology:** A quasi-experimental study with a non-equivalent control group and pretest-posttest measurements. Participants were 27 second-year Psychology students (74% female; mean age = 19 years). The experimental group (n=7) received three training sessions focused on Conscious Regulation of Behavior, Decision-Making, Emotional Regulation, and Monitoring of Responsibilities, identified through an initial assessment using the UEF-1 Scale. The control group (n=20) continued with their usual academic activities. Non-parametric analyses and the residual gain method were employed.

**Results:** The experimental group showed significant improvements in Conscious Regulation of Behavior (p = .026; r = .51), Emotional Regulation (p = .030; r = .49), and the Supervisory Attention System (p = .046; r = .44), with large effect sizes. The control group experienced no significant changes in any of the functions evaluated.

**Conclusions:** A brief, personalized program can enhance specific executive functions in university students, demonstrating cognitive plasticity in young adults. The findings support the design of contextually relevant interventions to strengthen transversal competencies in higher education.

## Introduction

Executive functions (EF) constitute a set of higher-order cognitive processes that enable self-regulation, planning, decision-making, and emotional control—fundamental skills for academic performance and social adaptation in adulthood [1]. In the university context, these abilities are particularly relevant, as higher education demands intensive and sustained use of these processes. Organizing independent study, regulating impulsivity in the face of academic demands, managing stress, and making decisions under pressure are daily challenges that can be compromised when executive functioning is not optimal [2].

However, the research tradition on the development and training of EFs has frequently focused on child populations, a stage in which they are considered especially plastic [3,4]; but studies focusing on young adults, particularly university students, are comparatively scarce. This imbalance in the literature is noteworthy, given that higher education demands intensive use of these capacities and support programs are often less structured than at previous educational levels [5]. The scarcity of research in this age group constitutes a significant knowledge gap, as we do not know to what extent brief, focused interventions can enhance these functions at a stage where cognitive plasticity still exists.

In the university population, seven key executive functions have been identified to date: the supervisory attention system, conscious regulation of behavior, conscious regulation of emotions, decision-making, conscious monitoring of responsibilities, management of elements to solve tasks, and verification of behavior for learning [6,7]. Furthermore, it has been verified that these executive functions are interrelated [8].

In the last decade, several studies have explored the efficacy of cognitive training programs to strengthen EFs in different age groups. Research such as that by Dunning et al. [9] has shown that interventions focused on self-control, mindfulness, and emotional regulation can generate significant improvements in children and adolescents. Likewise, work such as that by Kim et al. [10] has evidenced that brief meditation practices reduce distraction interference and optimize supervisory attention in adults. Furthermore, Díaz-Guerra et al. [11] point out the importance of integrating artificial intelligence-based academic tools as a resource for enhancing executive functions, considering the theory of distributed cognition. However, most of these programs have been designed with a generic approach, without considering the specific needs of the target population.

This methodological limitation has been highlighted by authors such as Yardley et al. [12], who propose a person-based approach for the development of interventions, adapting the content to the particular characteristics and deficits of the participants. In the university context, where academic demands are heterogeneous and executive development trajectories can vary considerably, personalization of programs emerges as a promising strategy to maximize their effectiveness [13]. Based on these postulates, this research aims to evaluate the effect of the University Executive Network-Training Program (NExT-U) on the improvement of executive functions in university students.

In this regard, the theoretical framework of this research is grounded in the explanatory model of executive functioning for university students by Díaz-Guerra et al. [14]. These authors demonstrate, through decision trees and structural equations, that executive functions in the university population do not operate as isolated processes, but are organized as a dynamic, stratified, and interdependent system. This reciprocal and multidirectional nature confirms that executive functions are organized as a network of influences where processes are mutually constructed and reinforced, providing a robust theoretical basis for the study of this population.

## Methodology

### Research Design

A quasi-experimental non-equivalent control group design with pretest and posttest measurements was employed. This methodological approach was selected because participant assignment to groups was not random but voluntary, a common situation in natural educational contexts where strict randomization is not feasible. With these designs, if the treatment is effective, post-test scores are expected to improve relative to pre-test scores [15].

In this design, the Independent Variable (IV) is the training program itself, while the Dependent Variable (DV) is the participants’ executive functioning. In the following representation, O1 represents the pre-test (baseline assessment), X represents the intervention, and O2 represents the post-test (assessment after the intervention) (see Figure 1).

**Figure 1.**
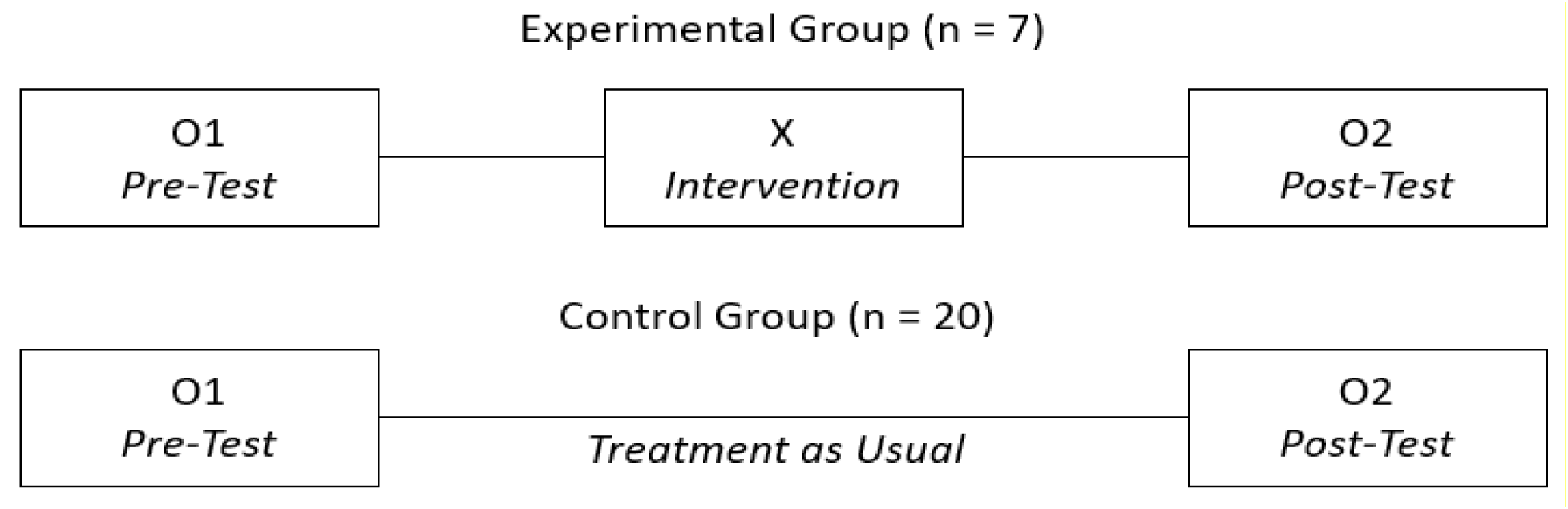
Pretest-posttest quasi-experimental design

### Participants

The research sample was non-probabilistic and intentional, consisting of 27 second-year participants enrolled in the Bachelor’s Degree in Psychology at the Central University “Marta Abreu” of Las Villas. Of these, 74% were female and the remaining 26% were male. The mean age was 19 years, ranging from 19 to 21 years. Additionally, the mean grade point average was 4.50.

Inclusion criteria were:

- Being officially enrolled in the second year of the Bachelor’s Degree in Psychology at the Central University “Marta Abreu” of Las Villas.
- Having availability to participate in all program sessions.
- Providing written informed consent.

The experimental group consisted of 7 students who voluntarily offered to participate in the program after the common pretest evaluation in both groups. The control group consisted of 20 students who, having completed the pretest, did not participate in the training program but continued with their academic activities. This group allowed for controlling maturation, history, and standardized testing effects.

Participants were prospectively recruited between October 20 and 24, 2025. Recruitment was conducted through announcements in classes and institutional communications at the University Well-being Center.

### Description of the University Executive Network-Training Program (NExT-U)

The University Executive Network-Training Program (NExT-U) is conceived as a specialized service integrated into the support system offered by the University Well-being Center [16] of the Central University “Marta Abreu” of Las Villas. Its theoretical foundation stems from an innovative conception of executive functions, understood as a dynamic system of interrelated and interdependent capacities that can be enhanced through structured and contextually relevant interventions [14].

One of the main strengths of NExT-U lies in its operational versatility, materialized in two implementation modalities designed to respond to different needs and institutional objectives. On the one hand, the Comprehensive Program Modality constitutes a systemic training itinerary aimed at the harmonious development of the entire spectrum of executive functions. This modality follows a fixed curricular progression and is aimed at students who, without presenting significant difficulties, aspire to reach higher levels of excellence in self-regulation and cognitive maturity. In this sense, it operates as a primary prevention and executive profile enrichment strategy, consolidating key transversal competencies for academic and professional success.

On the other hand, the Needs-Adapted Program Modality responds to a logic of diagnostic targeting that substantially distinguishes it from the previous one. Instead of covering all curricular content, this modality starts from an initial assessment that identifies the executive functions with the greatest difficulties in each student or group. Based on this diagnosis, specific intervention modules are selected, aimed at enhancing those areas that present functional vulnerability or priority needs. This flexibility allows maximizing intervention efficiency by concentrating resources on the dimensions that truly require support.

The program addresses the enhancement of the seven executive functions empirically identified in the university context: supervisory attention system, conscious regulation of behavior, conscious regulation of emotions, decision-making, conscious monitoring of responsibilities, management of elements to solve tasks, and verification of behavior for learning. In its comprehensive version, the intervention is structured over a seven-week cycle, with a frequency of three weekly sessions lasting one hour each, totaling twenty-one meetings that allow systematic and progressive work on each of the functions considered.

### Pre- and Post-Intervention Assessment Instruments

#### Executive Functions Scale for University Students (UEF-1)

This scale was designed and validated for Spanish-speaking participants by Ramos-Galarza et al. [6] in a sample of Chilean and Ecuadorian students and validated in the Cuban population by Díaz-Guerra et al. [7]. It consists of 31 items and measures 7 executive functions. The subscales it assesses are: Conscious Monitoring of Responsibilities (F1, items 2, 8, 9, 15, and 27), Supervisory Attention System (F2, items 10, 14, 22, 28, and 13), Conscious Regulation of Behavior (F3, items 3, 11, 16, 17, 18, and 19); Behavior Checking for Learning (F4, items 20, 23, 24, and 30); Decision Making (F5, items 5, 12, and 21), Conscious Regulation of Emotions (F6, items 4, 25, 29, and 31), and Management of Elements to Solve Tasks (F7, items 1, 6, 7, and 26). Responses are given on a 5-point Likert scale (1 = strongly disagree to 5 = strongly agree).

### Procedures and Data Analysis Strategy

The University Executive Network-Training Program (NExT-U) was applied at the University Well-being Center of the Central University “Marta Abreu” of Las Villas from November 3 to 5, 2025, each day at 2:00 pm. The pretest and posttest were administered one week before and after the program (respectively), at 10:00 am.

Based on the pretest results, Conscious Regulation of Behavior was identified as the executive function with the highest scores below average. Additionally, Decision Making, Conscious Regulation of Emotions, and Conscious Monitoring of Responsibilities were identified as executive functions susceptible to enhancement due to their scores, so they were secondarily integrated into the program. Based on this diagnosis, the Specific Program modality according to the sample’s needs was selected.

Control was maintained over the following variables, considered as confounding or extraneous variables to ensure group equivalence: sociodemographic data (age, gender, employment status), academic data (grade point average, hours of independent study per week), health and lifestyle variables (perceived sleep quality, sports habits, use of psycho-stimulant substances), prior experience (participation in similar workshops, courses, or therapies), use of academic artificial intelligence (prior experience and knowledge). Additionally, the presentation of the UEF-1 was randomized to avoid habituation bias in participants.

Statistical analyses were performed using Jamovi v.2.6 and G*Power 3.1 software. A 95% confidence level was used for all intervals and significance tests. Prior to inferential analyses, a data preparation and description stage was conducted. Subsequently, descriptive statistics were calculated to characterize the sample. For quantitative variables (UEF-1 sum scores, age, grade point average, weekly study hours), medians, means, and standard deviations were reported. For categorical variables (UEF-1 score ratings, gender, employment status, stimulant use, etc.), frequencies and percentages were calculated. These analyses were presented separately for the experimental group (n=7) and control group (n=20).

Considering the non-random nature of group assignment and the small sample size, non-parametric statistical tests were implemented to verify initial equivalence between groups at pretest. The Mann-Whitney U test for independent samples [17] was applied to each of the 7 UEF subscales at pretest, aiming to detect initial significant differences between groups. A significance level of α = .05 was used. To ensure comparability of groups on potentially confounding variables, comparisons were made between quantitative variables (age, grade point average, study hours) using the Mann-Whitney U test [17] and categorical variables (gender, concurrent work, sleep quality, stimulant use, prior experience with similar programs, AI knowledge and use) using Chi-Square and Fisher’s Exact tests [18].

To evaluate changes attributable to the training program and due to the sample size and violation of normality assumptions, the residual gain method was employed [19], using the control group in this case as a reference to establish the expected pretest-posttest trajectory in the absence of intervention. Under the assumption that residual gains represent the deviation of observed performance from that expected under natural conditions. The procedure was developed in three stages:

1. *Modeling the natural trajectory:* A linear regression was performed separately for each executive function, using exclusively data from the control group (n=20). In each model, the criterion variable was the posttest score and the predictor variable was the pretest score, thus establishing the expected trajectory without intervention.
2. *Prediction of expected values:* The regression values (constant and beta) obtained in each model were applied to all participants (experimental and control groups) to calculate the predicted posttest, representing the expected performance if they had not received the intervention. This was done using the formula: *Predicted_Posttest = Constant + (β x Observed_Pretest)*.
3. *Calculation of residual gains:* For each participant and executive function, the difference between the observed posttest and the predicted posttest was calculated using the formula: *Residual_Gain = Observed_Posttest – Predicted_Posttest*.

The residual gains for each executive function were compared between groups using the Mann-Whitney U test, evaluating whether the experimental group showed significantly greater improvements than the control group. To quantify the magnitude of the differences found, Rosenthal’s r effect size was calculated for each Mann-Whitney U comparison; values were interpreted according to conventional criteria: r = .10 (small effect), .30 (medium effect), .50 (large effect) [20]. A post-hoc power analysis was performed in G*Power v.3.1 software to determine the design’s capacity to detect existing effects.

### Ethical Considerations

At all stages of the research, participants were informed about the voluntary nature of their collaboration, as well as the absence of reprisals in case of withdrawal. Anonymity of responses and confidentiality of results were ensured. Informed consent was obtained from all adult participants included in the study. All procedures performed followed the ethical standards of the 1964 Declaration of Helsinki and its subsequent amendments, and comparable ethical standards [21]. The research protocol was approved by the Ethics Committee for Research with Humans of the Department of Psychology, Faculty of Social Sciences, Central University “Marta Abreu” of Las Villas (17OCT2025ID089).

## Results

### Descriptive Analysis and Data Preparation

The initial descriptive analysis allowed characterizing the participant profile and verifying the composition of the experimental and control groups regarding continuous control variables (see Table 1). Regarding age, both groups showed a similar age profile, with ages mainly in the 19-year-old range. Specifically, the control group had a mean of 19.50 years (SD = .69), while the experimental group showed a slightly lower mean of 19.29 years (SD = .49). Both groups shared the same median (19.00 years), suggesting a comparable age distribution between conditions.

**Table 1.**
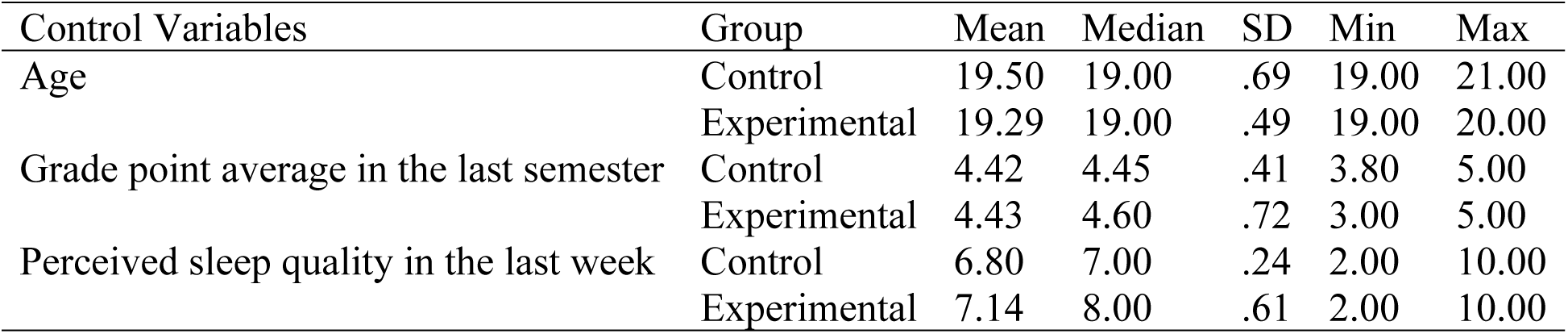
Descriptive statistics of continuous control variables.

| Control Variables | Group | Mean | Median | SD | Min | Max |
| --- | --- | --- | --- | --- | --- | --- |
| Age | Control | 19.50 | 19.00 | .69 | 19.00 | 21.00 |
|  | Experimental | 19.29 | 19.00 | .49 | 19.00 | 20.00 |
| Grade point average in the last semester | Control | 4.42 | 4.45 | .41 | 3.80 | 5.00 |
|  | Experimental | 4.43 | 4.60 | .72 | 3.00 | 5.00 |
| Perceived sleep quality in the last week | Control | 6.80 | 7.00 | .24 | 2.00 | 10.00 |
|  | Experimental | 7.14 | 8.00 | .61 | 2.00 | 10.00 |

Regarding prior academic performance, measured by the grade point average of the last semester, both groups maintained similarly high performance. The control group obtained a mean of 4.42 (SD = .41) with a median of 4.45, while the experimental group recorded a mean of 4.43 (SD = .72) and a median of 4.60. Although the means were practically identical, it is noteworthy that the experimental group showed greater variability in their grades, with a range spanning from 3.00 to 5.00, compared to the narrower range of the control group (3.80 to 5.00).

Regarding perceived sleep quality during the last week, assessed on a scale from 0 to 10, both groups reported moderately high levels. The control group had a mean of 6.80 (SD = 2.40) with a median of 7.00, while the experimental group showed slightly higher values, with a mean of 7.14 (SD = 2.61) and a median of 8.00. This difference, although small, might suggest a slightly more positive perception of sleep quality in the experimental group.

Regarding categorical control variables, these are presented in Table 2. Concerning gender, a marked female predominance was observed in both groups. Specifically, the control group consisted of 19 women (70%) and 1 man (4%), while the experimental group had 6 women (22%) and 1 man (4%). This distribution indicates a similar gender proportion between conditions, which minimizes the potential confounding effect of this variable.

**Table 2.**
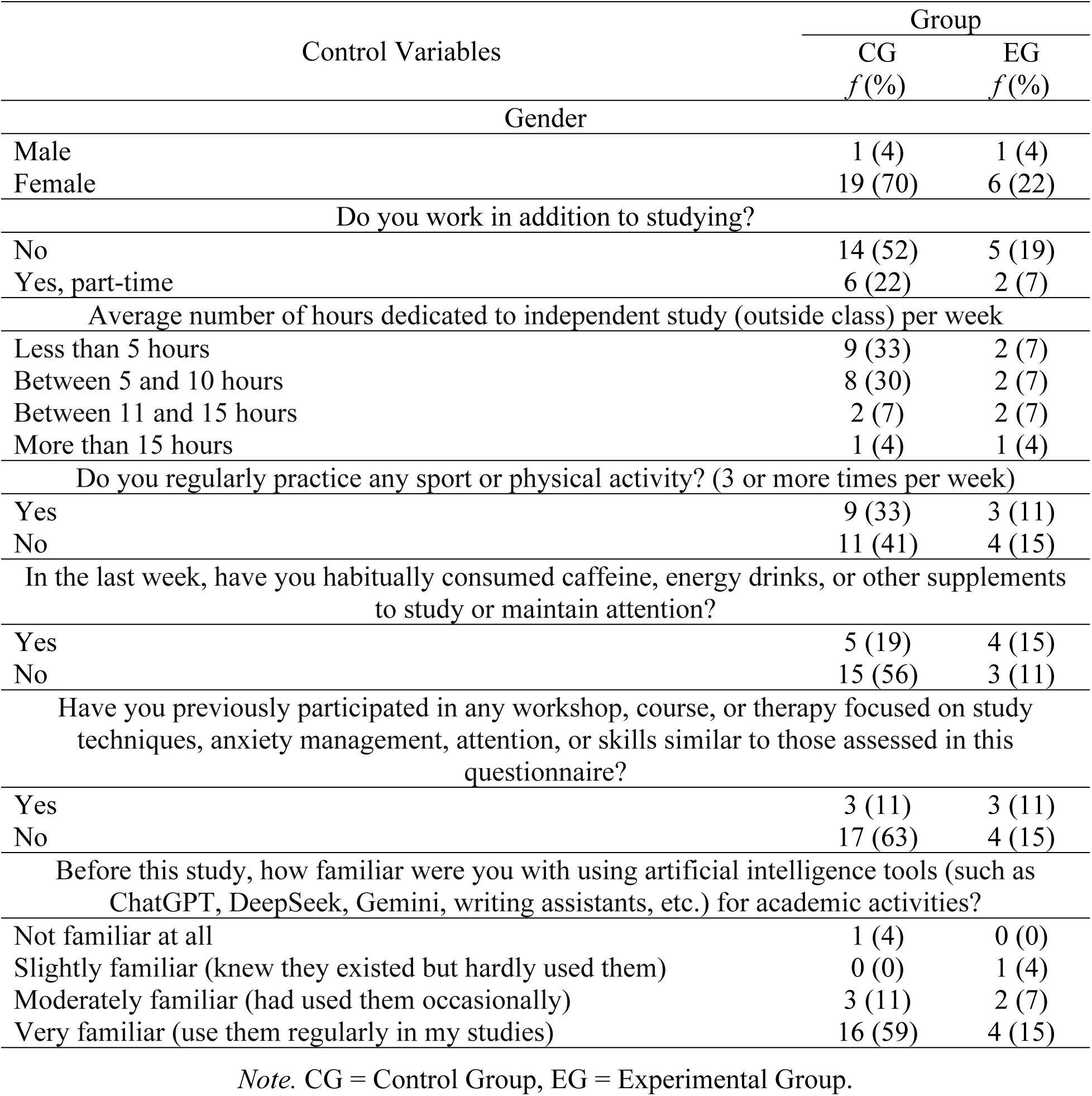
Descriptive statistics of categorical control variables.

| Control Variables | Group |  |
| --- | --- | --- |
|  | CG<br>*f* (%) | EG<br>*f* (%) |
| Gender |  |  |
| Male | 1 (4) | 1 (4) |
| Female | 19 (70) | 6 (22) |
| Do you work in addition to studying? |  |  |
| No | 14 (52) | 5 (19) |
| Yes, part-time | 6 (22) | 2 (7) |
| Average number of hours dedicated to independent study (outside class) per week |  |  |
| Less than 5 hours | 9 (33) | 2 (7) |
| Between 5 and 10 hours | 8 (30) | 2 (7) |
| Between 11 and 15 hours | 2 (7) | 2 (7) |
| More than 15 hours | 1 (4) | 1 (4) |
| Do you regularly practice any sport or physical activity? (3 or more times per week) |  |  |
| Yes | 9 (33) | 3 (11) |
| No | 11 (41) | 4 (15) |
| In the last week, have you habitually consumed caffeine, energy drinks, or other supplements to study or maintain attention? |  |  |
| Yes | 5 (19) | 4 (15) |
| No | 15 (56) | 3 (11) |
| Have you previously participated in any workshop, course, or therapy focused on study techniques, anxiety management, attention, or skills similar to those assessed in this questionnaire? |  |  |
| Yes | 3 (11) | 3 (11) |
| No | 17 (63) | 4 (15) |
| Before this study, how familiar were you with using artificial intelligence tools (such as ChatGPT, DeepSeek, Gemini, writing assistants, etc.) for academic activities? |  |  |
| Not familiar at all | 1 (4) | 0 (0) |
| Slightly familiar (knew they existed but hardly used them) | 0 (0) | 1 (4) |
| Moderately familiar (had used them occasionally) | 3 (11) | 2 (7) |
| Very familiar (use them regularly in my studies) | 16 (59) | 4 (15) |

*Note.* CG = Control Group, EG = Experimental Group.

Regarding employment status, most participants in both groups did not combine studies with paid work. In the control group, 14 students (52%) stated they did not work, and 6 (22%) worked part-time. A similar pattern was observed in the experimental group, where 5 participants (19%) did not work and 2 (7%) worked part-time. Regarding hours dedicated to weekly independent study, the distribution was varied, although with some concentration in the lower categories. In both the control group (33%) and the experimental group (7%), the “less than 5 hours” category was the most frequent, followed by “between 5 and 10 hours” (30% in control, 7% in experimental).

In the area of health and study habits, approximately one-third of each group practiced regular physical activity (33% in control, 11% in experimental). In contrast, stimulant use for studying showed a notable difference between groups: while in the control group only 5 participants (19%) reported habitual use, in the experimental group this figure was 4 participants (15%), representing a higher proportion within their group. Regarding prior experience in similar programs, both groups presented equivalent proportions, with 3 participants (11%) in each group reporting previous participation in workshops or courses focused on study techniques or anxiety management.

Finally, concerning familiarity with artificial intelligence tools for academic activities, both groups showed a considerable level of experience. The “very familiar” category was the most frequent in both cases, encompassing 16 participants (59%) in the control group and 4 (15%) in the experimental group. It is noteworthy that no participant fell into the “not familiar at all” category in the experimental group, while in the control group only 1 participant (4%) was in this category.

The descriptive analysis of executive functioning levels at pretest revealed a general pattern where most participants were located in the lower categories of the performance continuum (see Table 3). Specifically, for the seven executive functions evaluated, no participant in either group reached the categories of “above average” or “well above average” in the initial assessment. In the control group, a particularly concerning distribution was observed in conscious regulation of emotions (F6), where 12 participants (44%) were in the “well below average” category, while none were placed “below average.” A similar, though less marked, pattern was identified in conscious regulation of behavior (F3), with 14 participants (52%) in “below average” and 3 (11%) in “well below average.”

**Table 3.** Descriptive statistics of executive functioning levels at Pretest and Posttest.

| Factors |  | Well below average |  | Below average |  | Average |  | Above average |  | Well above average |  |
| --- | --- | --- | --- | --- | --- | --- | --- | --- | --- | --- | --- |
|  |  | *f*(%) |  | *f*(%) |  | *f*(%) |  | *f*(%) |  | *f*(%) |  |
|  |  | CG | EG | CG | EG | CG | EG | CG | EG | CG | EG |
| F1 | Pretest | 2 (7) | 1 (4) | 9 (33) | 2 (7) | 9 (33) | 4 (15) | 0 (0) | 0 (0) | 0 (0) | 0 (0) |
|  | Posttest | 2 (7) | 0 (0) | 9 (33) | 3 (11) | 9 (33) | 4 (11) | 0 (0) | 0 (0) | 0 (0) | 0 (0) |
| F2 | Pretest | 3 (11) | 0 (0) | 8 (30) | 2 (7) | 9 (33) | 5 (19) | 0 (0) | 0 (0) | 0 (0) | 0 (0) |
|  | Posttest | 5 (19) | 0 (0) | 4 (15) | 1 (4) | 11 (41) | 6 (22) | 0 (0) | 0 (0) | 0 (0) | 0 (0) |
| F3 | Pretest | 3 (11) | 0 (0) | 14 (52) | 5 (19) | 3 (11) | 2 (7) | 0 (0) | 0 (0) | 0 (0) | 0 (0) |
|  | Posttest | 4 (15) | 0 (0) | 12 (44) | 3 (11) | 4 (15) | 4 (15) | 0 (0) | 0 (0) | 0 (0) | 0 (0) |
| F4 | Pretest | 1 (4) | 1 (4) | 8 (30) | 1 (4) | 11 (41) | 5 (19) | 0 (0) | 0 (0) | 0 (0) | 0 (0) |
|  | Posttest | 2 (7) | 0 (0) | 4 (15) | 2 (7) | 14 (52) | 5 (19) | 0 (0) | 0 (0) | 0 (0) | 0 (0) |
| F5 | Pretest | 3 (11) | 0 (0) | 11 (41) | 4 (15) | 6 (22) | 3 (11) | 0 (0) | 0 (0) | 0 (0) | 0 (0) |
|  | Posttest | 4 (15) | 0 (0) | 7 (26) | 3 (11) | 9 (33) | 4 (15) | 0 (0) | 0 (0) | 0 (0) | 0 (0) |
| F6 | Pretest | 12 (44) | 2 (7) | 0 (0) | 0 (0) | 8 (30) | 5 (19) | 0 (0) | 0 (0) | 0 (0) | 0 (0) |
|  | Posttest | 7 (26) | 1 (4) | 0 (0) | 0 (0) | 13 (48) | 6 (22) | 0 (0) | 0 (0) | 0 (0) | 0 (0) |
| F7 | Pretest | 2 (7) | 0 (0) | 6 (22) | 0 (0) | 12 (44) | 7 (26) | 0 (0) | 0 (0) | 0 (0) | 0 (0) |
|  | Posttest | 1 (4) | 0 (0) | 6 (22) | 1 (4) | 13 (48) | 6 (22) | 0 (0) | 0 (0) | 0 (0) | 0 (0) |

*Note.* CG = Control Group, EG = Experimental Group, F1 = Conscious Monitoring of Responsibilities, F2 = Supervisory Attention System, F3 = Conscious Regulation of Behavior, F4 = Verification of Behavior for Learning, F5 = Decision Making, F6 = Conscious Regulation of Emotions, F7 = Management of Elements to Solve Tasks.

In contrast, the experimental group showed a slightly more favorable initial profile in several dimensions. For example, at pretest, no participant in the experimental group was placed in “well below average” for functions F2, F3, F5, and F7, while in F6 only 2 participants (7%) reached this lower category. However, as in the control group, no experimental participant exceeded the “average” category in any executive function during the initial assessment.

When comparing posttest distributions, differentiated patterns emerged between groups. The control group essentially maintained its distribution in most functions, with minor changes that did not substantially alter the overall picture. For example, in conscious monitoring of responsibilities (F1), the distribution remained identical between pretest and posttest, while in the supervisory attention system (F2), the number of participants in the “well below average” category increased from 3 to 5 (19%). However, some moderate improvements were observed in specific functions: in conscious regulation of emotions (F6), the percentage in “well below average” decreased from 44% to 26%, with a corresponding increase in the “average” category from 30% to 48%.

On the other hand, the experimental group showed a more positive evolution in several dimensions. Notably, at posttest, no experimental participant was placed in the “well below average” category for functions F1, F2, F3, F4, F6, and F7, completely eliminating this lower category in six of the seven functions evaluated. In verification of behavior for learning (F4), the number of participants in “well below average” decreased from 1 to 0, while in conscious regulation of emotions (F6), it decreased from 2 to 1 participant in this category. Furthermore, in all executive functions, the experimental group maintained or slightly increased its presence in the “average” category, consolidating this as their predominant location at posttest.

### Program Adjustment Based on Sample Needs

The descriptive analysis of pretest scores revealed that, within the experimental group, certain executive functions presented higher levels of difficulty before the implementation of the training program (see Figure 2). None of the executive functions had values above average or well above average.

**Figure 2.**
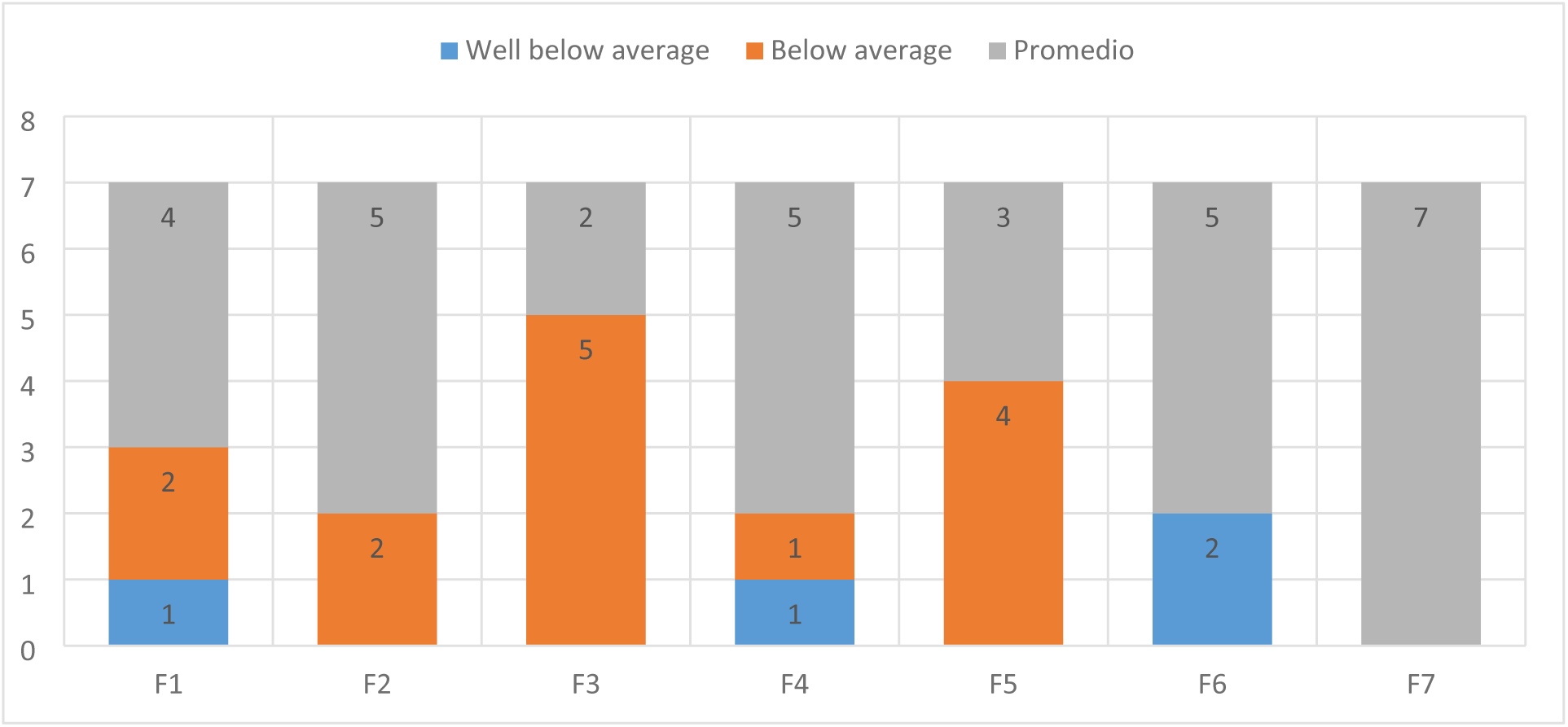
Levels of executive functioning in the experimental group pretest. *Note.* F1 = Conscious Monitoring of Responsibilities, F2 = Supervisory Attention System, F3 = Conscious Regulation of Behavior, F4 = Verification of Behavior for Learning, F5 = Decision Making, F6 = Conscious Regulation of Emotions, F7 = Management of Elements to Solve Tasks.

Specifically, Conscious Regulation of Behavior (F3) emerged as the most affected function, with 19% of participants (5 out of 7) located in categories below average. This indicates significant challenges in the ability to intentionally monitor and adjust behaviors.

Secondly, Decision Making (F5) also showed notable difficulties, with 15% of participants (4 out of 7) positioned below average, suggesting limitations in the processes of evaluating, selecting, and implementing options. Complementarily, Conscious Monitoring of Responsibilities (F1) presented 11% of participants (3 out of 7) in these lower categories, reflecting possible deficiencies in tracking and fulfilling academic and personal obligations.

Finally, Conscious Regulation of Emotions (F6) showed 7% of participants (2 out of 7) with performance well below average, evidencing challenges in the ability to manage emotional responses adaptively. Although the Supervisory Attention System (F2) and Verification of Behavior for Learning (F4) had the same scores as F6 (both with 2 participants total in the below average or well below average categories), F6 was identified as more susceptible to enhancement because all its values were located in the well below average category. This was not the case for F2, where all were in the below average category, nor for F3, which had one case in each category.

Based on the identification of these executive functions susceptible to enhancement, and considering the explanatory model of Conscious Regulation of Behavior, developed in Stage 2 of this research, the training program was directed toward these variables. In this sense, the sessions were restructured using the NExT-U modality of Specific Program Based on Sample Needs across 3 sessions (see Table 4).

**Table 4.** Description of the implemented sessions.

| Phase | Technique | Primary Executive Function | Secondary Executive Functions | Objective | Activity |
| --- | --- | --- | --- | --- | --- |
| Session 1: Self-Control and Reflective Pause Techniques |  |  |  |  |  |
| Opening | "The Mental Stop Game" | Conscious Regulation of Behavior | Decision Making (evaluation of options during the pause) | Identify behavioral automatisms and emotional state. | Students walk freely; upon saying "Stop!", they analyze: "What thought/action did I interrupt? What emotion did I feel?" + Record on an emotional thermometer. |
| Development | The 10-Second Pause Technique |  | Conscious Emotional Regulation (recognition of emotions during the "stop") | Break cycles of impulsivity by integrating emotional analysis. | Guided practice: Faced with a stimulus (e.g., mobile notification), wait 10 seconds before acting. Record: Note what they felt during the wait using an emotional format. |
|  | Behavioral Traffic Light with P.A.U.S.E. |  | Conscious Monitoring of Responsibilities (planning the "pause commitment") | Visualize levels of control and decision options. | Create a personal poster with: Red: "Stop me" + breathe 4-7-8 Yellow: "Reflect" using P.A.U.S.E. questions Green: "Act" consciously |
| Closing | "My SMART Pause Commitment" |  |  | Transfer learning to real situations with planning. | Each student defines: "I will apply the pause when [situation] to achieve [SMART goal]". E.g., "Before responding in class, I will breathe for 10 seconds to participate 3 times this week." |
| Session 2: Decision Making Under Pressure |  |  |  |  |  |
| Opening | "The Hourglass with Stress Traffic Light" | Conscious Regulation of Behavior | Decision Making (P.A.U.S.E. method and time-limited simulation) | Experience time-limited stress and identify emotional activation. | Solve a math puzzle in 2 minutes using color cards (red/yellow/green) to indicate stress level during the task. |
| Development | "Pop Quiz" Simulation |  | Conscious Emotional Regulation (4-7-8 breathing and anxiety management) | Manage anxiety in academic contexts with integrated techniques. | Scenario: "You have 15 minutes to solve an unannounced puzzle." Roles: Student using P.A.U.S.E. + 4-7-8 breathing vs. one who acts impulsively. |
|  | P.A.U.S.E. Method with Eisenhower Matrix |  | Conscious Monitoring of Responsibilities (self-assessment with "Productivity Thermometer") | Structure decisions under pressure with prioritization. | Steps: 1. Pause (4-7-8 breathing) 2. Analyze options using urgent/important matrix 3. Select choosing the best 4. Act 5. Review results |
| Closing | "My Decision Logbook" |  |  | Self-assess decision quality and progress. | Complete a record: 1 impulsive decision (red) + what to learn from it 1 thoughtful decision (green) + what made it effective Adjustment plan for the week |

| Session 3: Self-Regulation and Routines with AI |  |  |  |  |  |
| --- | --- | --- | --- | --- | --- |
| Opening | "My Day in Emojis with Coping Table" | Conscious Regulation of Behavior | Conscious Monitoring of Responsibilities (routines with AI and visual reminders) | Identify disorganization patterns and coping strategies. | Draw on a timeline how their previous day was using only emojis (e.g., 📱😓💻) and classify the strategies used as healthy/unhealthy. |
| Development | DeepSeek: Integrated Routine Assistant |  | Conscious Emotional Regulation ("Celebration of Micro-achievements" and validation) | Automate daily planning with emotional management. | Example command: "Create a 4-hour study routine after school, with breaks, stress management, and time for social media." Include in the routine: Emotional emergency kit and progress check-ins. |
|  | Visual Reminders with Bias Checklist |  | Decision Making (using AI to foresee consequences of routines) | Externalize self-control and avoid distractions. | Design cards with icons to paste in strategic places: 🚫📱 (no cell phone) + "Is it urgent or can I wait?" ⌚ (time to start) + "Breathe 4-7-8 before beginning" |
| Closing | "Letter to My Future Self with Celebration of Micro-achievements" |  |  | Consolidate learning and reinforce self-regulated behaviors. | Each student writes a letter with: 1 new strategy they will use + 1 warning about their "usual trap." They share 1 micro-achievement and the group responds "Way to go!" + snap. |

### Verification of Group Equivalence at Pretest

The analysis of normality assumptions, using the Shapiro-Wilk test, revealed a non-normal distribution in most of the evaluated control variables, thus justifying the choice of non-parametric statistical tests for subsequent analyses (see Table 5). Of the ten variables analyzed, nine presented significant violations of the normality assumption (p < 0.05), indicating that their distributions deviated substantially from a normal curve. Specifically, variables such as age (p < .001), gender (p < .001), employment status (p < .001), grade point average (p = .011), hours of independent study (p = .020), physical activity practice (p < .001), stimulant use (p < .001), prior experience in similar programs (p < .001), and familiarity with AI tools (p < .001) showed non-parametric distributions according to established statistical criteria.

**Table 5.**
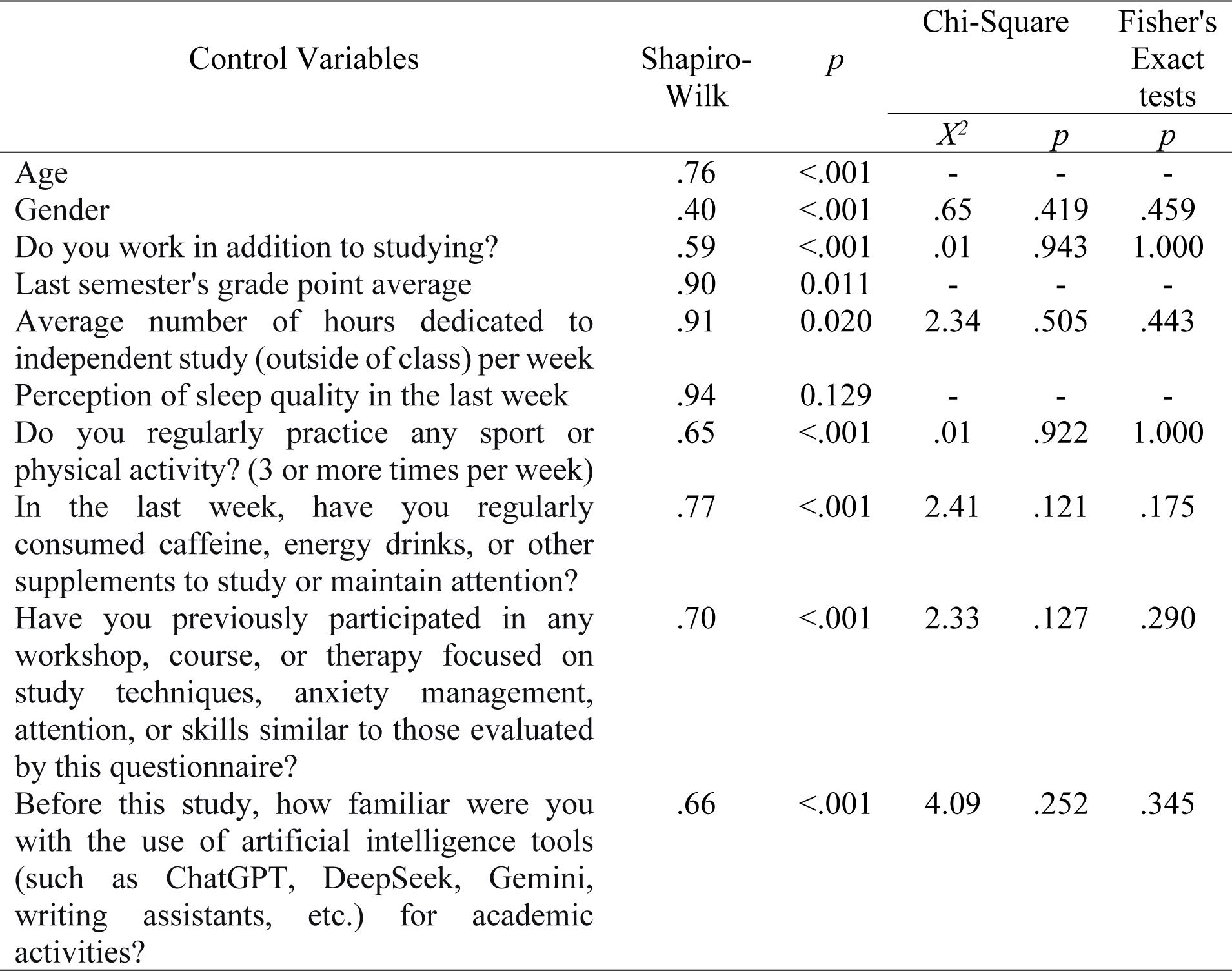
Evaluation of normality assumptions and equivalence between groups.

| Control Variables | Shapiro-Wilk | $p$ | Chi-Square | | Fisher's Exact tests |
| --- | --- | --- | --- | --- | --- |
| | | | $X^2$ | $p$ | $p$ |
| Age | .76 | <.001 | - | - | - |
| Gender | .40 | <.001 | .65 | .419 | .459 |
| Do you work in addition to studying? | .59 | <.001 | .01 | .943 | 1.000 |
| Last semester's grade point average | .90 | 0.011 | - | - | - |
| Average number of hours dedicated to independent study (outside of class) per week | .91 | 0.020 | 2.34 | .505 | .443 |
| Perception of sleep quality in the last week | .94 | 0.129 | - | - | - |
| Do you regularly practice any sport or physical activity? (3 or more times per week) | .65 | <.001 | .01 | .922 | 1.000 |
| In the last week, have you regularly consumed caffeine, energy drinks, or other supplements to study or maintain attention? | .77 | <.001 | 2.41 | .121 | .175 |
| Have you previously participated in any workshop, course, or therapy focused on study techniques, anxiety management, attention, or skills similar to those evaluated by this questionnaire? | .70 | <.001 | 2.33 | .127 | .290 |
| Before this study, how familiar were you with the use of artificial intelligence tools (such as ChatGPT, DeepSeek, Gemini, writing assistants, etc.) for academic activities? | .66 | <.001 | 4.09 | .252 | .345 |

Only the variable of perceived sleep quality in the last week (p = .129) did not reach statistical significance in the Shapiro-Wilk test, suggesting that its distribution might approximate normality. However, considering that the vast majority of variables presented clear violations of the normality assumption, and given the reduced sample size (N = 27) which limits the robustness of parametric tests, a conservative approach was chosen by applying non-parametric methods to all variables to maintain methodological consistency in the analyses.

Subsequently, an initial equivalence analysis was performed using non-parametric Mann-Whitney U tests. In this analysis, it can be observed that there are no significant differences between the experimental and control groups in any of the continuous variables evaluated at pretest. For the age variable, no significant differences were found between groups (U = 60.00, p = .537), indicating that the age distribution is equivalent. Regarding the grade point average of the last semester, the analysis showed equivalence between groups (U = 58.00, p = .518), demonstrating that prior academic performance was comparable. In perceived sleep quality, no significant differences were found either (U = 60.00, p = .594), confirming similar levels between groups.

The verification of initial equivalence between groups in categorical variables, using Chi-square and Fisher’s Exact tests, revealed the absence of statistically significant differences in all evaluated dimensions, thus confirming the adequate comparability of the groups before the implementation of the training program (see Table 5). Specifically, regarding basic demographic characteristics, no significant differences were found in gender distribution (p = .459 in Fisher’s exact test) nor in the employment status of participants (p = 1.000), indicating that both groups had similar compositions in these fundamental aspects.

In relation to study habits and well-being, the comparisons showed statistical equivalence in all analyzed variables. Regarding hours dedicated to weekly independent study (p = .443), regular practice of physical activity (p = 1.000), and stimulant use to maintain attention (p = .175), both groups exhibited comparable distributions, suggesting similar academic behavior and health patterns before the intervention. This similarity is particularly relevant, as variables such as study time and stimulant use could potentially affect performance in executive functions.

Finally, regarding participants’ prior experience, the analyses confirmed that there were no significant differences between groups either in previous participation in similar programs (p = .290) or in the level of familiarity with artificial intelligence tools for academic activities (p = .345). This equivalence in prior experience is methodologically crucial, as it ensures that any subsequent differences in results cannot be attributed to baseline advantages in previous knowledge or training.

The analysis of normality assumptions for pretest scores in the seven evaluated executive functions, using the Shapiro-Wilk test, evidenced significant violations of the normal distribution assumption in all evaluated dimensions (see Table 6). Specifically, each of the executive functions showed p-values below the established significance level (p = 0.05), thus confirming non-parametric distributions in the initial assessment. In particular, functions F1: Conscious monitoring of responsibilities (p < .001), F3: Conscious regulation of behavior (p < .001), F4: Verification of behavior for learning (p < .001), F6: Conscious regulation of emotions (p < .001), and F7: Management of elements to solve tasks (p < .001) presented the most marked violations of normality. Complementarily, although with slightly less extreme significance levels, but still below the critical threshold, functions F2: Supervisory attention system (p = 0.002) and F5: Decision making (p = 0.002) also showed distributions that deviated significantly from the normal curve.

**Table 6.** Evaluation of normality assumptions and equivalence of executive functions at pretest.

| Factors | Shapiro-Wilk | *p* | Mann-Whitney U | *p* |
| --- | --- | --- | --- | --- |
| F1: Conscious monitoring of responsibilities | .79 | <.001 | 64.000 | .737 |
| F2: Supervisory attention system | .86 | 0.002 | 48.500 | .197 |
| F3: Conscious regulation of behavior | .84 | <.001 | 53.000 | .256 |
| F4: Behavior checking for learning | .74 | <.001 | 62.000 | .633 |
| F5: Decision making | .87 | 0.002 | 55.000 | .367 |
| F6: Conscious regulation of emotions | .79 | <.001 | 48.000 | .170 |
| F7: Management of elements to solve tasks | .79 | <.001 | 42.000 | .057 |

The evaluation of initial equivalence between the experimental and control groups in pretest executive function scores, using the Mann-Whitney U test, revealed the absence of statistically significant differences in all evaluated dimensions, thus confirming the adequate comparability of the groups before the implementation of the intervention program (see Table 6). Specifically, in the seven executive functions analyzed, none of the p-values reached the established significance level (p = 0.05), evidencing similar distributions between groups at baseline assessment. The function showing the smallest difference between groups was F1: Conscious monitoring of responsibilities (U = 64.000, p = .737), followed very closely by F4: Verification of behavior for learning (U = 62.000, p = .633) and F5: Decision making (U = 55.000, p = .367).

Consistently, the remaining functions also showed statistical equivalence between groups, although with slightly lower p-values that did not reach significance. Particularly, F3: Conscious regulation of behavior (U = 53.000, p = .256), F2: Supervisory attention system (U = 48.500, p = .197), and F6: Conscious regulation of emotions (U = 48.000, p = .170) presented probability levels that remained considerably above the significance threshold. It is worth noting that function F7: Management of elements to solve tasks (U = 42.000, p = .057) showed the p-value closest to the critical level, although without reaching statistical significance, suggesting a marginal difference that, nevertheless, does not compromise the overall equivalence of the groups according to conventional statistical criteria.

This uniform pattern of non-significance in all comparisons provides solid evidence that the experimental and control groups were statistically equivalent in their baseline levels of executive functioning before the intervention. The consistency of these results considerably strengthens the internal validity of the quasi-experimental design, as it minimizes the probability that any difference subsequently observed in the posttest could be attributed to initial disparities in the participants’ executive capacities. This baseline equivalence constitutes a robust methodological foundation for attributing subsequent changes to the effect of the implemented training program.

### Evaluation of the Program Effect

The evaluation of intra-group changes between pretest and posttest, using the Wilcoxon W test for related samples, revealed differentiated patterns of evolution in executive functions between the experimental group and the control group (see Table 7). In the experimental group, statistically significant improvements were observed in three of the seven executive functions evaluated after the implementation of the training program. Specifically, the function that showed the most notable improvement was F3: Conscious regulation of behavior (W = .00, p = .017), followed by F2: Supervisory attention system (W = .00, p = .049) and F6: Conscious regulation of emotions (W = .00, p = .016). These significant improvements were supported by large effect sizes (r = −1.00 in all three functions), indicating substantial and consistent changes in these dimensions.

**Table 7.**
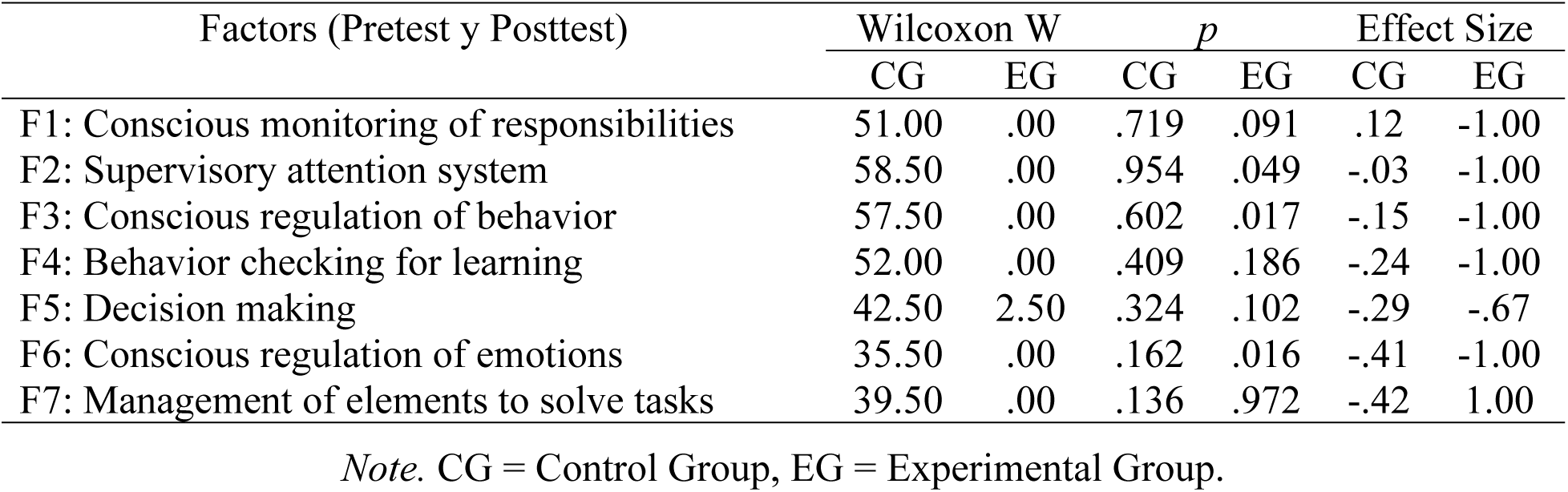
Evaluation of intra-group changes between pretest and posttest.

Complementarily, in the experimental group, improvement trends that did not reach conventional statistical significance but merit consideration were identified. Functions F1: Conscious monitoring of responsibilities (p = .091) and F5: Decision making (p = .102) presented p-values close to the significance threshold, with effect sizes of −1.00 and −.67 respectively, suggesting possible improvements that could reach statistical significance with greater sample power. It is particularly interesting to observe that function F7: Management of elements to solve tasks (p = .972) showed a very high p-value accompanied by a positive effect size (r = 1.00), which could indicate that this function was already a strength of the experimental group and did not experience significant changes after the intervention.

In marked contrast, the control group showed no statistically significant improvements in any of the seven executive functions evaluated, with all p-values well above the established significance level (p = 0.05). Specifically, functions F1: Conscious monitoring of responsibilities (p = .719), F2: Supervisory attention system (p = .954), F3: Conscious regulation of behavior (p = .602), F4: Verification of behavior for learning (p = .409), F5: Decision making (p = .324), F6: Conscious regulation of emotions (p = .162), and F7: Management of elements to solve tasks (p = .136) maintained stable scores over time, with effect sizes ranging from small to moderate in negative or positive directions, but without reaching statistical significance. This stability in the control group constitutes an important reference point, as it demonstrates that, in the absence of intervention, executive functioning levels did not experience significant changes during the study period.

The construction of linear regression models, based exclusively on control group data, allowed establishing the expected change trajectories for each executive function in the absence of intervention, thus providing a baseline against which to evaluate the effectiveness of the training program (see Table 8). The analyses revealed that pretest was a significant predictor of posttest in all evaluated executive functions, with beta coefficients (β) ranging from .68 to 1.00, indicating consistent positive relationships between initial and final assessments. Specifically, function F1: Conscious monitoring of responsibilities showed a perfect relationship (β = 1.00) accompanied by a constant close to zero (.03), suggesting that in this dimension the posttest tended to replicate almost exactly the pretest values under non-intervention conditions.

**Table 8.**
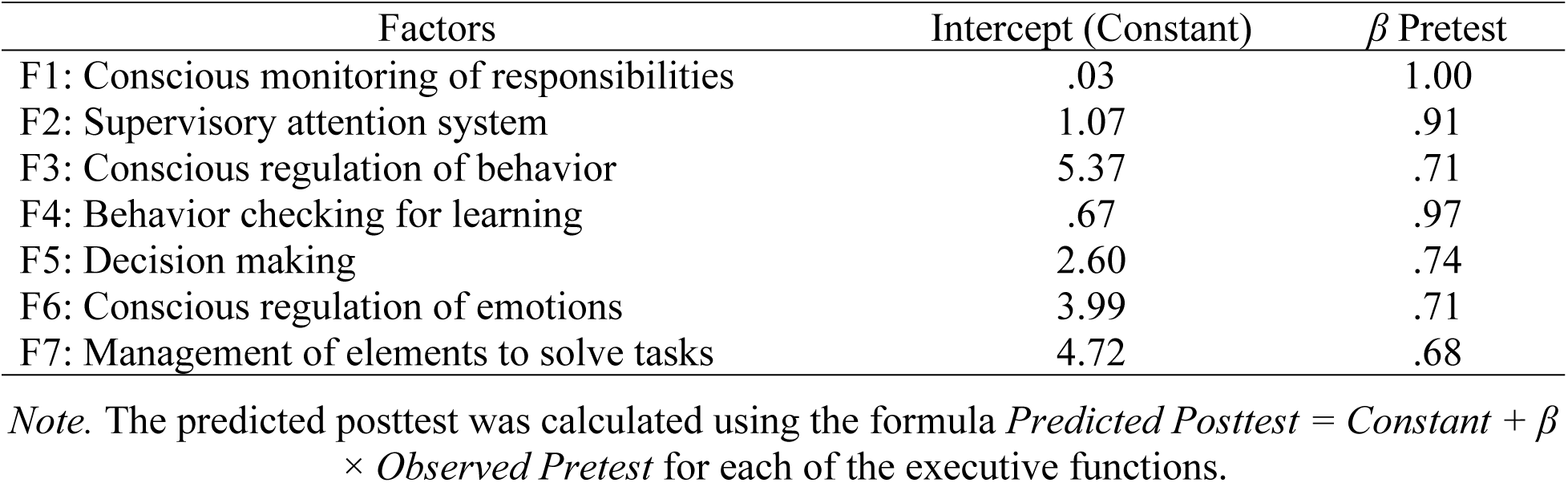
Constant values and betas of the linear regression models for calculating expected posttest based on pretest in the control group.

Complementarily, functions F2: Supervisory attention system (β = .91) and F4: Verification of behavior for learning (β = .97) presented particularly high coefficients, indicating that approximately 91% and 97% of the variability in the posttest could be explained by the initial pretest scores in these dimensions. In contrast, functions F7: Management of elements to solve tasks (β = .68), F3: Conscious regulation of behavior (β = .71), F6: Conscious regulation of emotions (β = .71), and F5: Decision making (β = .74) showed moderate beta coefficients, suggesting that, although pretest was a significant predictor of posttest, there was greater variability not explained by the initial assessment in these functions.

Based on these data, the residual gain was calculated using the formula Residual Gain = Observed Posttest – Predicted Posttest. The residual gain represents how much the participant’s actual posttest score deviates from the score that would have been expected without intervention, considering their baseline score. This calculation revealed differentiated patterns between the experimental and control groups (see Figure 3).

**Figure 3.**
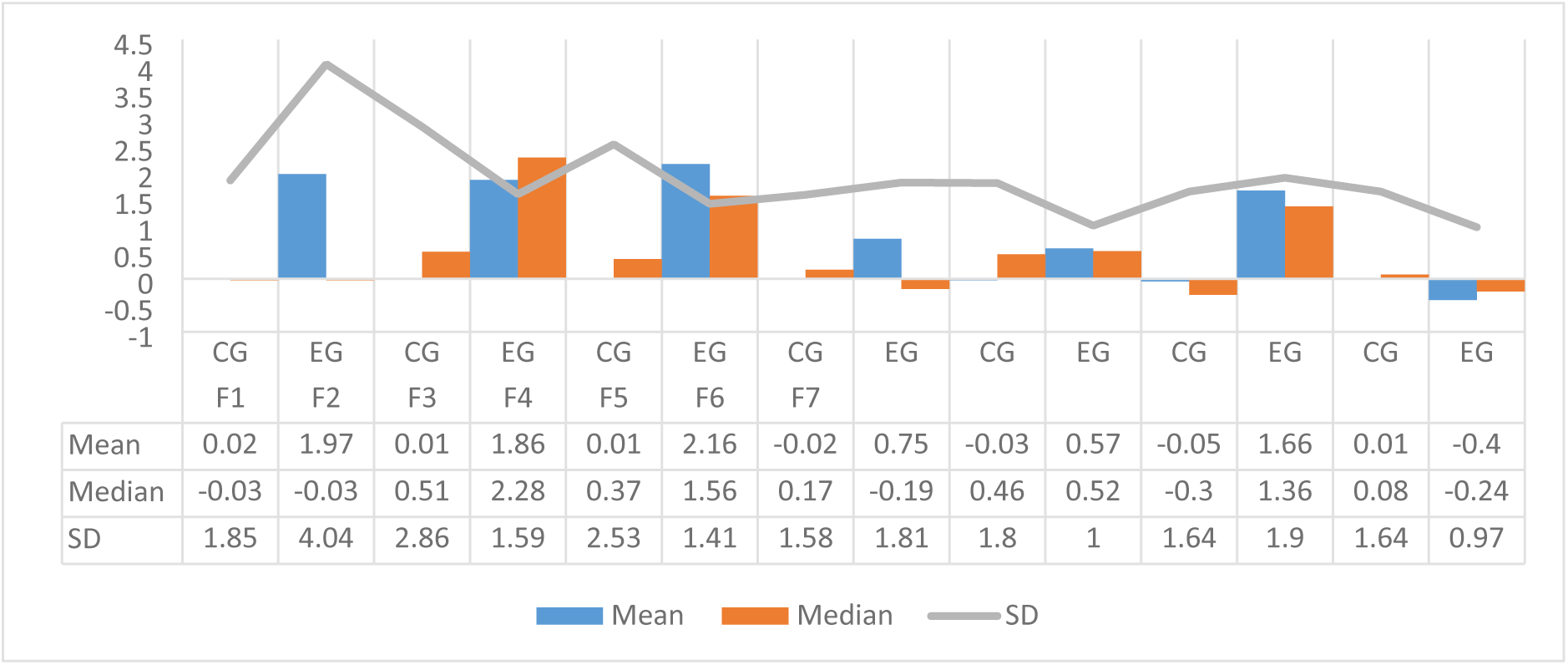
Residual gains in the control and experimental groups *Note*. CG = Control Group, EG = Experimental Group, F1 = Conscious Monitoring of Responsibilities, F2 = Supervisory Attention System, F3 = Conscious Regulation of Behavior, F4 = Verification of Behavior for Learning, F5 = Decision Making, F6 = Conscious Regulation of Emotions, F7 = Management of Elements to Solve Tasks.

In the experimental group, substantial positive residual gains were observed in six of the seven functions, particularly highlighting function F3: Conscious regulation of behavior with a mean of 2.16 (SD = 1.41) and median of 1.56, followed by F1: Conscious monitoring of responsibilities with a mean of 1.97 (SD = 4.04) and F2: Supervisory attention system with a mean of 1.86 (SD = 1.59). These positive gains indicate that participants in the experimental group consistently exceeded expectations based on their initial performance, achieving posttest scores significantly higher than what would have been predicted had they followed the natural trajectory of the control group.

Complementarily, the experimental group showed moderate but positive gains in functions F6: Conscious regulation of emotions (mean = 1.66, SD = 1.90), F4: Verification of behavior for learning (mean = 0.75, SD = 1.81), and F5: Decision making (mean = 0.57, SD = 1.00). It is particularly revealing that the only function where the experimental group showed a negative residual gain was F7: Management of elements to solve tasks (mean = −0.40, SD = 0.97), suggesting that, in this specific dimension, the experimental group’s posttest performance was slightly lower than what would have been expected based on their initial score and the control group’s trajectory.

In marked contrast, the control group presented residual gains close to zero in all executive functions, with means ranging from −0.05 to 0.02, confirming that their change trajectories closely aligned with those predicted by the regression models. Standard deviations in the control group were generally larger than in the experimental group for several functions (e.g., 2.86 in F2 vs. 1.59 in the experimental group), indicating greater variability in residual gains among participants. This variability, however, was distributed symmetrically around zero, as evidenced by the fact that the minimum and maximum values in the control group showed both residual gains and losses (e.g., −6.90 to 4.47 in F2), while in the experimental group residual gains tended to concentrate on positive values in most functions.

The comparison of residual gains between the experimental group and the control group, using the Mann-Whitney U test, revealed statistically significant differences in three of the seven executive functions evaluated. This constitutes evidence of differentiated effects of the training program on specific dimensions of executive functioning (see Figure 4).

**Figure 4.**
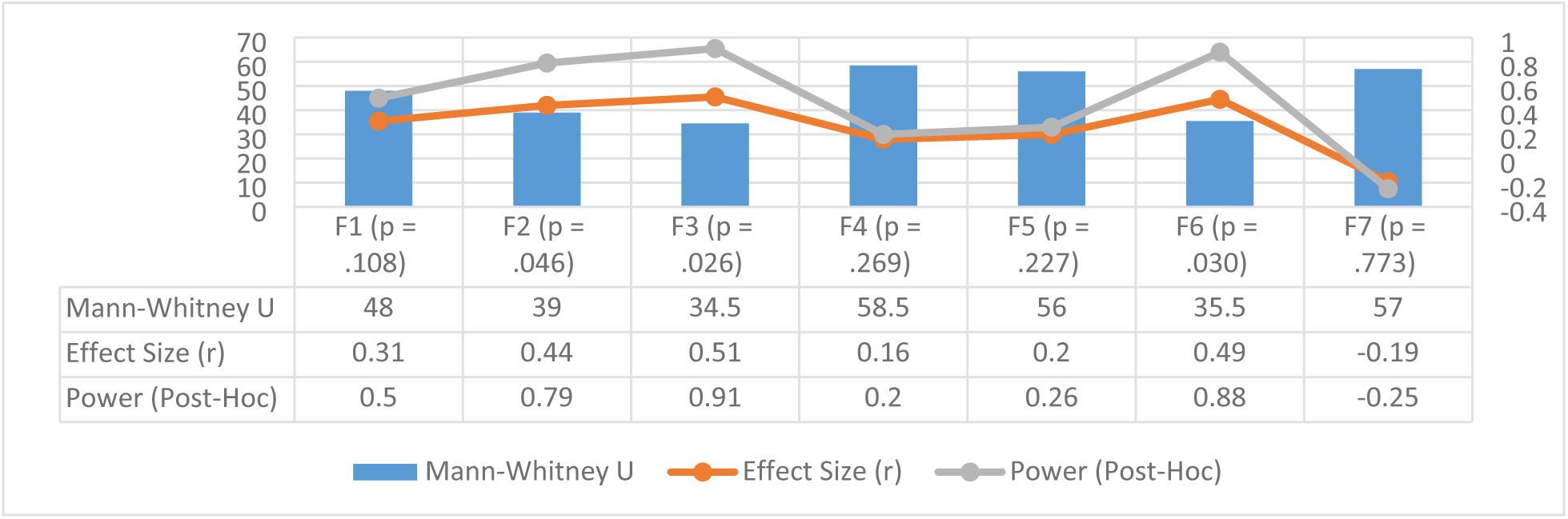
Comparison of effect size and power of residual gains in both groups *Note.* F1 = Conscious Monitoring of Responsibilities, F2 = Supervisory Attention System, F3 = Conscious Regulation of Behavior, F4 = Verification of Behavior for Learning, F5 = Decision Making, F6 = Conscious Regulation of Emotions, F7 = Management of Elements to Solve Tasks.

Particularly, the function showing the most marked difference was F3: Conscious regulation of behavior (U = 34.50, p = .026), with a large effect size (r = .51) and a high post-hoc statistical power of 91%, indicating that the training program generated substantial and statistically detectable improvements in this dimension. Similarly, function F6: Conscious regulation of emotions (U = 35.50, p = .030) also showed significant differences between groups, with a large effect size (r = .49) and power of 88%, confirming the program’s effectiveness in enhancing emotional regulation capacities. Complementarily, function F2: Supervisory attention system (U = 39.00, p = .046) reached statistical significance with a moderate-large effect size (r = .44) and power of 79%, suggesting that the program also positively impacted participants’ attentional mechanisms.

In contrast, four executive functions did not show statistically significant differences between groups, although they presented differentiated patterns in their results. Function F1: Conscious monitoring of responsibilities (p = .108) showed a moderate effect size (r = .31) and power of 50%, indicating an improvement trend that did not reach conventional statistical significance but could represent a real effect that the current design lacked sufficient power to detect. Similarly, functions F5: Decision making (p = .227, r = .20) and F4: Verification of behavior for learning (p = .269, r = .16) presented small effect sizes and low power (26% and 20% respectively), suggesting that the program had a limited impact on these specific dimensions.

It is particularly interesting that function F7: Management of elements to solve tasks showed not only a non-significant difference (p = .773), but also a negative effect size (r = −.19) and negative power, which coincides with previous findings that this function was already a relative strength of the experimental group at pretest and did not experience additional improvements after the intervention. The post-hoc power analyses revealed that the study had variable capacity to detect effects, ranging from 20% for small effects to 91% for large effects, providing valuable information for the design of future studies with larger sample sizes that could detect subtler effects in those functions where non-significant improvement trends were observed.

These results provide robust evidence that the training program was effective in specifically enhancing executive functions related to behavioral regulation, emotional regulation, and supervisory attention. While it showed limited or non-significant effects on functions related to decision-making, behavioral verification, and management of elements, particularly in those dimensions where the group already presented adequate performance in the initial assessment.

## Discussion

The results demonstrated that the implementation of the University Executive Network-Training Program (NExT-U), in its needs-based program modality, produced significant improvements in key executive functions among university students, particularly in Conscious Regulation of Behavior, Conscious Regulation of Emotions, and the Supervisory Attention System. These improvements were not only significant but also showed large effect sizes (r = .51, .49, and .44, respectively), suggesting considerable practical relevance. In contrast, the control group experienced no significant changes in any of the functions evaluated, which reinforces the hypothesis that the improvements observed in the experimental group are attributable to the intervention program.

These findings align with previous research highlighting the plasticity of executive functions in young adults and the efficacy of structured interventions to strengthen them. For instance, studies such as those by Diamond [22] and Salem et al. [23] have underscored that training in self-control and reflective pause techniques —similar to those applied in this program— can favor behavioral and emotional regulation, especially in academic contexts where pressure and impulsivity are frequent. Likewise, the improvement in the Supervisory Attention System coincides with reports by Fujino et al. [24], who found that integrating attentional focusing and distraction management techniques increases the capacity to monitor and adjust behavior in real time.

It is important to highlight that the program was specifically designed to primarily enhance Conscious Regulation of Behavior and, secondarily, Decision Making, Conscious Regulation of Emotions, and Conscious Monitoring of Responsibilities. The results confirm that significant improvements were achieved in the two primary functions that were the central focus of the intervention: Conscious Regulation of Behavior and Conscious Regulation of Emotions. Although it was not a direct objective, the Supervisory Attention System also showed significant improvement. This finding supports the systemic and comprehensive nature of executive functions, where intervening on a specific component can generate positive effects in other interconnected domains, thereby confirming in practice the results of the model by Díaz-Guerra et al. [14] regarding the bidirectionality and systemic correlation that characterizes the executive network. This interdependence has been noted by authors such as Friedman and Miyake [25] and Guo et al. [26], whose unity and diversity model proposes that, while executive functions are distinguishable, they operate interactively and cohesively, so that enhancing one can facilitate improvement in others, even without direct training.

Specifically, this improvement could be due to a cognitive prerequisite effect, given that attentional monitoring and detection mechanisms constitute the necessary substrate for any intentional regulation process [27]. Neurocognitively, this transfer can be understood through the overlap of prefrontal neural networks that support inhibitory control and supervisory attentional processes, according to Chidharom et al. [28].

A relevant aspect of this study was the design of the program based on an initial needs assessment [12], which allowed prioritizing the functions most susceptible to improvement. This personalized approach may explain why functions such as Conscious Regulation of Behavior and Emotional Regulation showed notable advances, in contrast to others like Conscious Monitoring of Responsibilities and Decision Making, which did not register significant changes. This suggests that, although executive functions are interdependent, their enhancement through brief interventions may be selective and depend on focusing on specific mechanisms [29], such as the inhibition of impulsive responses and the management of emotional states.

The absence of statistical significance in Conscious Monitoring of Responsibilities and Decision Making, despite having been addressed in the program, can be interpreted by considering several interrelated factors. From a neurocognitive perspective, these functions represent higher-order metacognitive processes that might require longer consolidation periods than those provided in a brief intervention. This is consistent with the functional hierarchy proposed by Díaz-Guerra et al. [14], according to which more complex functions are built upon the basis of more elementary processes and, therefore, could manifest changes later or require prior consolidation of their prerequisites. This is corroborated by authors such as Kälin & Roebers [30], who suggest that inhibition functions, such as Conscious Regulation of Emotions and Conscious Regulation of Behavior, effectively constitute necessary prerequisites for the optimal development of complex metacognitive capacities.

It is worth noting that, although the experimental group showed positive residual gains in most functions, the heterogeneity in the results —especially in Conscious Monitoring of Responsibilities— indicates that not all participants responded to the program in the same way. This could be related to individual differences in motivation, learning styles, or level of adherence to the proposed strategies, a phenomenon also observed in studies such as that by Sala et al. [31] on cognitive training transfer.

The methodological limitations of the study offer a complementary explanation for these results. The small size of the experimental group (n = 7) may have provided insufficient statistical power to detect small or moderate effect magnitudes, particularly in those functions where improvement trends were observed that did not reach conventional significance (Conscious Monitoring of Responsibilities: p = .108; Decision Making: p = .102). Therefore, the selective effectiveness of the program —significant in regulatory functions, but not in metacognitive ones— should not be interpreted as a limitation, but rather as a contribution to the understanding of change mechanisms in executive training. The results support the view of executive functions as a dynamic system where different components respond to intervention at different times and magnitudes, depending on their complexity, their interconnection with other functions, and their sensitivity to contextual characteristics [14,32,33].

From an applied perspective, these results have relevant implications for the design of support programs for university students, especially in contexts where dropout and low performance are associated with difficulties in self-regulation and emotional management [34]. The incorporation of practical sessions, based on simulation and using tools such as artificial intelligence for planning, appears to be a promising way to strengthen key transversal competencies for academic and professional success. However, it is important to acknowledge some limitations, such as the small sample size —especially in the experimental group— and the lack of long-term follow-up, which prevents determining whether the effects are maintained over time.

## Conclusions

The results of this study demonstrate that a brief three-session training program, designed based on an initial needs assessment, produces significant improvements in specific executive functions of university students. Positive effects with large effect sizes were observed in Conscious Regulation of Behavior, Conscious Regulation of Emotions, and the Supervisory Attention System, confirming that young adults maintain sufficient cognitive plasticity to benefit from focused interventions.

The selective effectiveness of the program (significant in regulatory functions but not in those of a complex metacognitive nature such as Decision Making) supports theoretical models that conceive executive functions as a dynamic and interdependent system. The improvement in the Supervisory Attention System, despite not being a direct target, evidences transfer relationships between interconnected executive components, consistent with the explanatory model of Díaz-Guerra et al. [14].

The stability of the control group in all evaluated functions reinforces the internal validity of the findings, allowing the improvements to be attributed to the implemented program and not to maturational or practice factors. Likewise, the person-based approach to intervention design proved to be a valid strategy to maximize the relevance and effectiveness of the program. Among the study’s limitations, the small sample size stands out (especially in the experimental group, n=7), which limited the statistical power to detect small effects, and the absence of long-term follow-up measurements to assess the sustainability of the changes. Future research should replicate these findings with larger samples and incorporate delayed assessments. On an applied level, these results suggest that brief, practical, and contextually relevant interventions can constitute an efficient complement to university support services, contributing to strengthening key transversal competencies for academic and professional success.

## Declarations

### Ethics approval and consent to participate

The research protocol was reviewed and approved by the Institutional Ethics Committee of the Department of Psychology, Faculty of Social Sciences, Universidad Central “Marta Abreu” de Las Villas (Approval No. 17OCT2025ID089).

### Consent for publication

Written informed consent for the publication of data from this study was obtained from all participants. Participants were informed that the data would be published anonymously and that non-essential identifying details would be omitted.

### Availability of data and materials

The datasets generated and analyzed during this study are available from the corresponding author upon reasonable request.

## Author Contribution Statements

- Diego D. Díaz Guerra: Conceptualization, Data Curation, Formal Analysis, Investigation, Methodology, Project Administration, Resources, Software, Supervision, Validation, Visualization, Writing – Original Draft, Writing – Review & Editing.
- Evelyn Fernández Castillo: Conceptualization, Formal Analysis, Investigation, Methodology, Resources, Validation, Visualization, Writing – Original Draft, Writing – Review & Editing.
- Carlos Ramos Galarza: Conceptualization, Formal Analysis, Investigation, Methodology, Resources, Validation, Visualization, Writing – Original Draft, Writing – Review & Editing.
- Melany de la Torre Pérez: Formal Analysis, Investigation, Methodology, Resources, Software, Validation, Visualization, Writing – Original Draft, Writing – Review & Editing.
- Yelenys González Espinosa: Formal Analysis, Investigation, Methodology, Resources, Software, Validation, Visualization, Writing – Original Draft, Writing – Review & Editing.
- Marena de la C. Hernández Lugo: Conceptualization, Formal Analysis, Investigation, Methodology, Resources, Validation, Visualization, Writing – Original Draft, Writing – Review & Editing.
- Vania Lugones Dapresa: Formal Analysis, Investigation, Methodology, Resources, Validation, Visualization, Writing – Original Draft, Writing – Review & Editing.
- Yunier Broche Pérez: Formal Analysis, Investigation, Methodology, Resources, Validation, Visualization, Writing – Original Draft, Writing – Review & Editing.

## Acknowledgments

This research was carried out as part of Diego D. Díaz Guerra’s Master’s studies in Clinical and Educational Neuropsychology and Doctoral studies in Psychological Sciences.

